# Inducible expression of the restriction enzyme uncovered genome-wide distribution and dynamic behavior of histones H4K16ac and H2A.Z at DNA double-strand breaks in *Arabidopsis*

**DOI:** 10.1101/2023.03.09.531848

**Authors:** Kohei Kawaguchi, Mei Kazama, Takayuki Hata, Mitsuhiro Matsuo, Junichi Obokata, Soichirou Satoh

## Abstract

DNA double-strand breaks (DSBs) are among the most serious types of DNA damage and cause mutations and chromosomal rearrangements. In eukaryotes, DSBs are immediately repaired in coordination with chromatin remodeling for the deposition of DSB-related histone modifications and variants. To elucidate the details of DSB-dependent chromatin remodeling throughout the genome, artificial DSBs need to be reproducibly induced at various genomic loci. Recently, in mammals, a comprehensive method for elucidating chromatin remodeling at multiple DSB loci via chemically induced expression of a restriction enzyme was developed. However, this DSB induction system is not suitable for investigating chromatin remodeling during and after DSB repair, and such approach has not been performed in plants. Here, we established a transgenic *Arabidopsis* plant harboring a restriction enzyme gene *Sbf* I driven by a heat-inducible promoter. Using this transgenic plant, we performed chromatin immunoprecipitation followed by deep sequencing (ChIP-seq) of histones H4K16ac and H2A.Z and investigated dynamics of these histone marks around the endogenous 623 *Sbf* I recognition sites. We also precisely quantified DSB efficiency at all cleavage sites using the DNA resequencing data obtained by ChIP-seq procedures. From the results, *Sbf* I-induced DSBs were detected at 360 loci and induced the transient deposition of H4K16ac and H2A.Z around these regions. Interestingly, we also observed the co-localization of H4K16ac and H2A.Z at some DSB loci. Overall, DSB-dependent chromatin remodeling was found to be a highly conserved between plants and animals. These findings provide new insights into chromatin remodeling that occurs in response to DSBs in *Arabidopsis*.

## Introduction

Organisms are constantly exposed to the risk of DNA damage by ultraviolet rays, ionizing radiation, and reactive oxygen species. DNA double-strand breaks (DSBs) are among the most serious types of DNA damage and can cause mutations and chromosomal rearrangements (Antunes et al. 2012, Shen et al. 2017, Varga and Aplan 2005). To maintain genomic integrity, both prokaryotes and eukaryotes, including plants and algae, have developed sophisticated DSB repair mechanisms. In eukaryotes, DSBs on the chromosomes are repaired in coordination with chromatin remodeling for the deposition of DSB-related histone modifications and variants (Clouaire and Legube 2019, Price and D’Andrea 2013). Such chromatin remodeling during DSB repair is required to facilitate the binding of DSB repair factors to the DSB loci and subsequent restoration of the DNA ends (Price and D’Andrea 2013, Lukas et al. 2011, Shi and Oberdoerffer 2012, Van and Santos 2018). In particular, the alterations of histone modifications and variants regulate DNA accessibility; promote the cohesiveness, flexibility, and mobility of chromatin within the nucleus; and facilitate the recruitment of specific factors ensuring repair and its related transcription. Recently, key factors for the interaction between DSB repair mechanisms and the chromatin landscape have gradually been uncovered. Chromatin remodelers such as SWI/SNF complexes, Tip60, histone acetyltransferases (HATs), and histone deacetylases (HDACs) tightly control the acetylation levels of histone H3 and H4 tails to regulate chromatin relaxation in the vicinity of DSBs (Lee et al. 2010, Murr et al. 2006, Miller et al. 2010, Dhar et al. 2017). H4K16ac, H3K14ac, and H3K56ac play particularly important roles in DNA damage response, open chromatin formation, and repair factor recruitment (Li et al. 2010, Kim et al. 2009, Chen et al. 2008, Aricthota et al. 2022). In addition, the NuA4-Tip60 complex introduces H2A.Z, a variant of histone H2A, onto nucleosomes near the DSBs and promotes acetylation of histone H4 (Gursoy-Yuzugullu et al. 2015, Xu et al. 2012). H2A.Z accumulation after DNA damage is transient and rapidly removed by Anp32e in mammalian cells (Gursoy-Yuzugullu et al. 2015, Obri et al. 2014), but the counterpart (including homolog of Anp32e) in plants is unclear. Altogether, these specific chromatin marks not only establish open chromatin states for efficient DNA repair, but also directly contribute to activation of the DNA damage response, the recruitment of DSB repair machinery, and the selection of repair pathway. Some histone marks have been reported to be deposited during DSB repair in plants (Lang et al. 2012, Hirakawa et al. 2019), but data on such chromatin remodeling are extremely limited, and the elucidation of such mechanisms has lagged considerably behind equivalent mammalian research. Therefore, a comprehensive survey of the patterns and frequency of chromatin remodeling during DSB repair in plants is crucial to obtain a better understanding of them.

To perform the genome-wide analysis to elucidate the details of DSB-induced chromatin remodeling, artificial DSBs need to be reproducibly induced at various genomic loci. Genotoxic agents (e.g., bleomycin, zeocin) and ionizing radiation, often used to induce DSBs, generate random breaks throughout the genome (thus heterogeneously across the cell population), and thus they are unsuitable for reproducible chromatin analyses. Meanwhile, DSB induction systems such as ZFN (Kim et al. 1996), TALEN (Christian et al. 2010), and CRISPR-Cas9 (Mali et al. 2013, Cong et al. 2013) induce only one localized DSB, which is insufficient for comprehensive analysis. Recently, genome-wide and site-directed induction of DSBs followed by ChIP-Seq analysis was performed and elucidated chromatin remodeling across the entire mammalian genome during the DSB repair process (Iacovoni et al. 2010, Iannelli et al. 2017, Clouaire et al. 2018). This DSB induction system is a very powerful tool to comprehensively investigate local chromatin remodeling at multiple DSB loci; however, such DNA cleavage system uses the chemical inducer, and thus DNA cleavage enzymes continue to be produced sustainably until such inducers are removed from cells. Thus, this system is not ideal for tracking chromatin dynamics after DSB repair.

Here, we established a transgenic *Arabidopsis* plant harboring a restriction enzyme gene *Sbf* I driven by heat-inducible promoter. This DNA cleavage enzyme recognizes an 8 bp specific sequence at the endogenous 623 loci in the *Arabidopsis* nuclear genome. Using this transgenic plant, we performed a comprehensive analysis of the distribution patterns of histones H4K16ac and H2A.Z around *Sbf* I recognition sites during and after DSB induction. We also precisely quantified DSB efficiency at all cleavage sites by using the DNA resequencing data obtained by ChIP-seq procedures. The results revealed DSBs at 360 loci, and we then observed the localization of H4K16ac and H2A.Z in many of these regions. Our method makes it possible to combine high-resolution mapping of DSBs with ChIP-seq analyses for DSB-related histone marks and provide detailed profiles of the dynamics of DNA repair and chromatin remodeling in the plant genome.

## Results

### Heat shock induction of the restriction enzyme gene *Sbf* I in *Arabidopsis thaliana*

DNA double-strand breaks (DSBs) need to be induced at a large number of targeted loci throughout the plant genome, and followed by estimation of DSB efficiency for their loci. To establish a model system to examine a DSB-related chromatin remodeling in plants, we introduced a gene encoding the restriction endonuclease *Sbf* I into the *Arabidopsis* genome. The recognition sequence of this enzyme is CCTGCA/GG, which is sparsely distributed at the endogenous 623 loci in the *Arabidopsis* nuclear genome. This number of recognition sites is smaller than for other restriction enzymes, and thus physiological damage by the overaccumulation of *Sbf* I-dependent DSBs is expected to be relatively limited (Supplementary Fig. S1A, Supplementary Table S1). In addition, we adopted the *Arabidopsis* heat-inducible promoter HSP18.2 for transcription of the *Sbf* I gene (Fig. 1A), and showed that the transcription level of the introduced *Sbf* I gene is elevated up to approximately 6,000-fold after heat shock (HS) treatment at 37°C for 4 hours, which then rapidly returns to its basal level over 2 additional days (Fig. 1B, C, Supplementary Fig. S1B, C). These results indicate that these *Sbf* I-inducible plants could easily and quickly regulate *Sbf* I gene expression simply by HS treatment. On the basis of these results, we used the transgenic line *Sbf* I-7 for subsequent analyses.

**Fig. 1.**
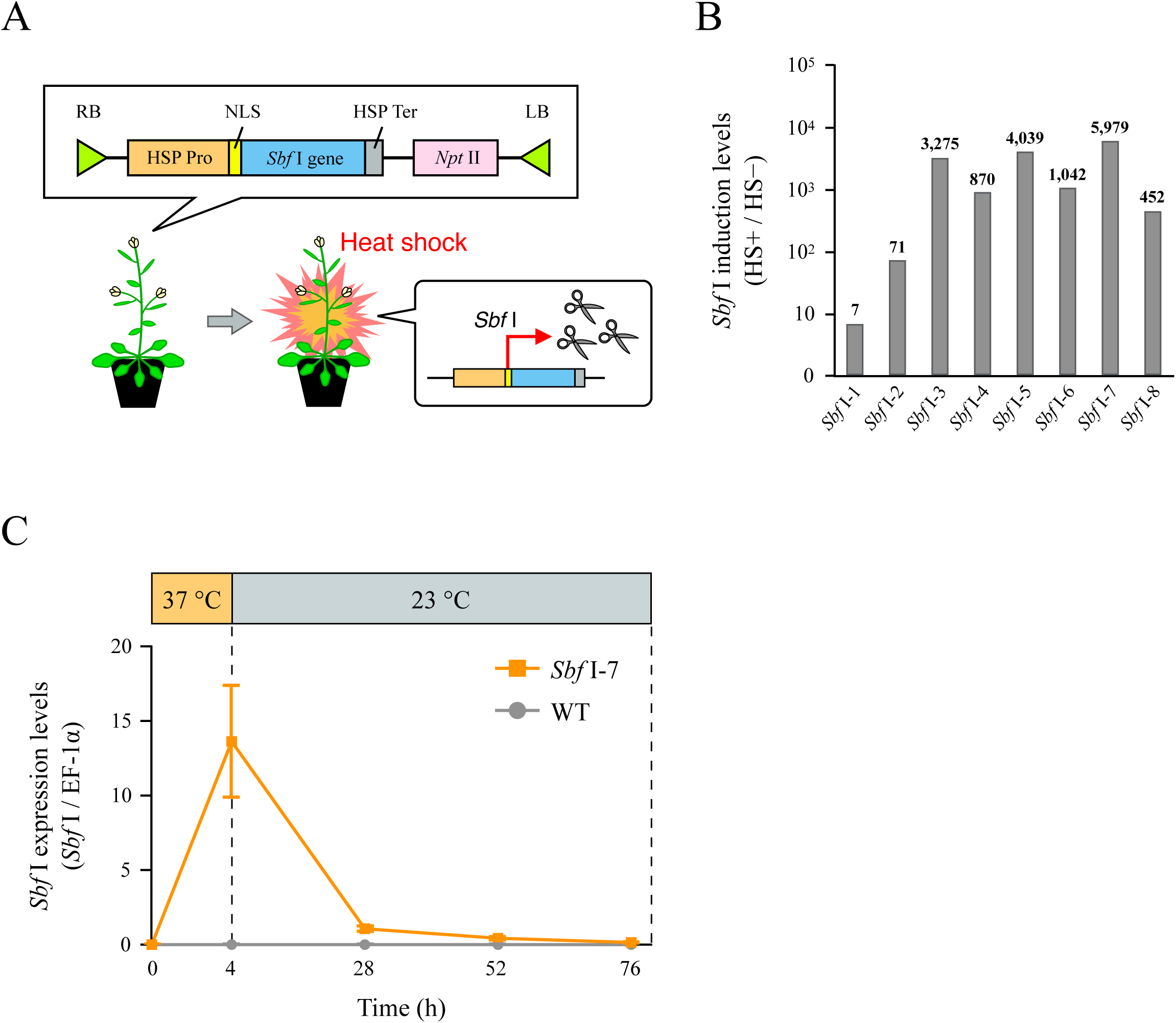
Establishment of heat-inducible *Sbf* I expression system in *Arabidopsis thaliana*. (A) Outline of T-DNA construct introduced into *Arabidopsis thaliana* and *Sbf* I gene expression model by heat shock (HS) treatment. (B) Degrees of heat-induced transcription levels of the *Sbf* I gene by RT-qPCR in T1 plants. The values on the vertical axis show the intensity of heat-induced *Sbf* I expression levels after HS treatment (HS+) relative to that before HS treatment (HS-). Most transgenic lines significantly increased *Sbf* I transcription levels after HS treatment. In particular, the *Sbf* I-7 line showed no leaky *Sbf* I transcription before HS treatment, and had the highest heat-induced transcription level of approximately 6,000-fold (Supplementary Fig. S1B, C). (C) Transcription level of the *Sbf* I gene after HS treatment quantified by RT-qPCR. WT sample was used as a negative control. Error bars are ±SD of three biological replicates.

### Genome-wide analysis of *Sbf* I-mediated cleavage activity in *Arabidopsis* nuclear genome

Generally, chromatin immunoprecipitation (ChIP) experiments are performed by chromatin DNA, whereas DSBs are detected by a different DNA sample. To investigate DSB-induced chromatin remodeling more precisely, it is appropriate to perform DSB detection and ChIP experiment under the same condition. In this study, we performed next-generation sequencing (NGS) analysis using sheared chromosomal DNA for the ChIP analysis (referred to as input DNA) from *Sbf* I-7 plants to quantify the DSB efficiency at the *Sbf* I recognition sites (*Sbf* I RS) in the *Arabidopsis* genome. If DSBs occurred at the *Sbf* I RS upon HS treatment, DNA fragments that include the *Sbf* I RS would be shortened or degraded and difficult to detect by NGS analysis. Thus, genomic regions with lower sequence coverage around the *Sbf* I RS in the HS-treated samples should reflect the cleavage of DNA by *Sbf* I. Furthermore, owing to this scheme, we could detect DSB sites from the ChIP-seq sequencing data, because ChIP sample and input sample (ChIP negative control) are both stemmed from the same chromatin DNA. We thus compared the mapped reads of input DNA from before and after HS treatment in the regions 1 kb upstream and downstream of all *Sbf* I RS in the *Arabidopsis* genome. As shown in Fig. 2A, the coverages of 100 bp around the *Sbf* I RS were substantially decreased in the HS samples at 360 of the 623 *Sbf* I RS. The amount of digested DNA differed depending on the cleavage site; approximately 40% of the DNA was digested at the loci where DSB efficiency was the highest (Supplementary Fig. S2A, Supplementary Table S2). In the samples of 48 hours after HS treatment (the 52h samples), such low-coverage profile disappeared; thus indicating the successful artificial cleavage and subsequent DNA-repair (Fig. 2A, B). In contrast, wild-type (WT) plants did not show a decrease in coverage at 360 DSB loci after HS treatment (Fig. 2A, Supplementary Fig. S2B, C).

**Fig. 2.**
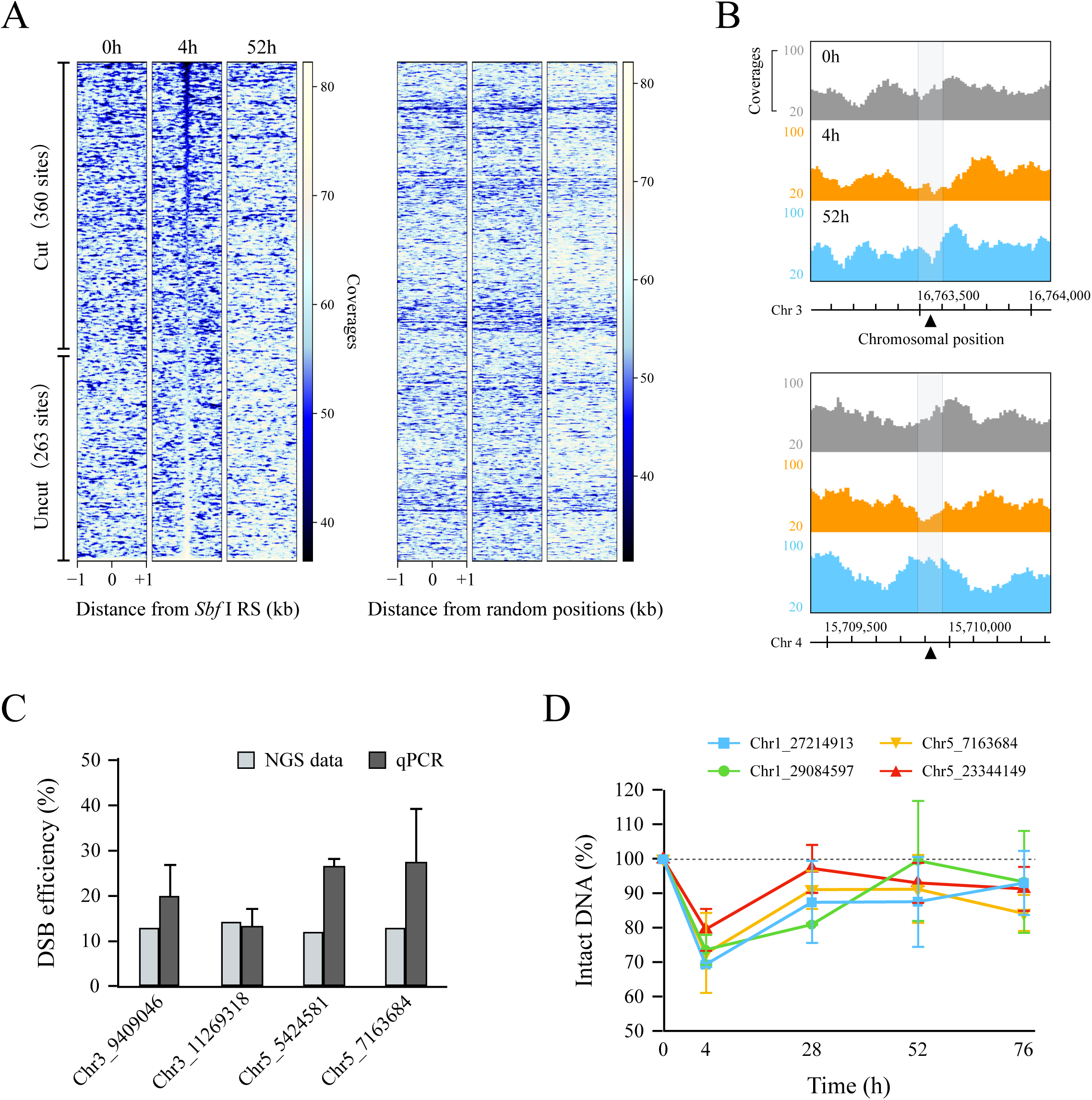
Quantification of DSB efficiency at 623 *Sbf* I recognition sites in the *Arabidopsis* nuclear genome. (A) Heatmap obtained from NGS data of input DNA in *Sbf* I-7 plants. These figures show the coverages around a 2 kb window surrounding the 623 *Sbf* I recognition sites (*Sbf* I RS) and 1,000 random positions. Left heatmaps were sorted in ascending order of the total coverages of 100 bp around the *Sbf* I RS for the 4h sample. (B) Genome browser screenshots representing the coverages of input DNA around the two *Sbf* I RS (chromosomes 3 and 4). The gray area covers 100 bp around DSB loci (black triangle). (C) Comparison of the DSB efficiency between NGS data and qPCR. The values of NGS data (gray blocks) and qPCR (dark gray blocks) were calculated by the quotient of the total coverages of 100 bp around the *Sbf* I RS (4h divided 0h) and by the difference of the PCR-amplified values (0h minus 4h), respectively. (D) The percentage of intact DNA by qPCR analysis at different time points. Each value was quantified relative to the amount of intact DNA before HS treatment set as 100%. Error bars are ±SD of three biological replicates.

To validate our quantification of the DSB efficiency from NGS data of input DNA, we performed qPCR analysis using primer sets designed to span each *Sbf* I RS, as shown in Supplementary Table S8. In principle, the difference of amplification efficiency by PCR between HS-treated and HS-untreated samples can be regarded as DSB efficiency. At the four *Sbf* I RS, DSB efficiencies quantified by qPCR were equal to or greater than those obtained from NGS analysis (Fig. 2C). Previously, Kozak et al. (2009) reported that most DNA ends were restored within hours to days when multiple DSBs were induced in the *Arabidopsis* genome. We also observed that DSBs at several cleavage sites were repaired up to 48 hours after HS treatment; two and four sites were repaired by 24 and 48 hours, respectively (Fig. 2D). These results indicated that, in most cells, DSBs were completely repaired within 48 hours after their occurrence. Hence, we demonstrated that *Sbf* I-inducible plants could simultaneously cause transient DSBs at various loci in the *Arabidopsis* genome, and are useful for comprehensive chromatin analysis during and after the DSB repair process. Hereafter, 360 loci where DSBs were observed were considered for ChIP analysis.

### DNA cleavage activity by *Sbf* I did not change in a locus-dependent manner

DNA cleavage enzymes are substantially influenced by the genetic context and the higher-order chromatin structures at chromosomal positions (Chung et al. 2020, Chen et al. 2016). To examine the relationship between DSB efficiency and the genetic context around the *Sbf* I RS, 360 DSB loci were classified into four groups (Gene, Intergenic, Transposable element, and Others) based on the TAIR10 genomic annotations (https://www.arabidopsis.org), and those classified into the Gene group were further divided into four groups (5′ and 3′ Untranslated regions, Exon, and Intron). Although we examined the relationship between DSB efficiencies and genomic context, we could not find any significances. (Fig. 3A-C). We next queried whether the three-dimensional chromatin structure interfere the DSB by *Sbf* I, particularly focused on the accessibility of DNA using PlantCADB (Ding et al. 2022), which included the data from various NGS analyses as follows: ATAC-seq, DNase-seq, FAIRE-seq, and MNase-seq. However, *Sbf* I-based DNA cleavage was also not significantly affected by the higher-order epigenetic context of the target sequences (Fig. 3A, D). Therefore, DSB efficiency by *Sbf* I does not depend on the DNA accessibility, and these results strongly suggest that *Sbf* I-inducible plants are versatile DSB induction tools *in vivo*.

**Fig. 3.**
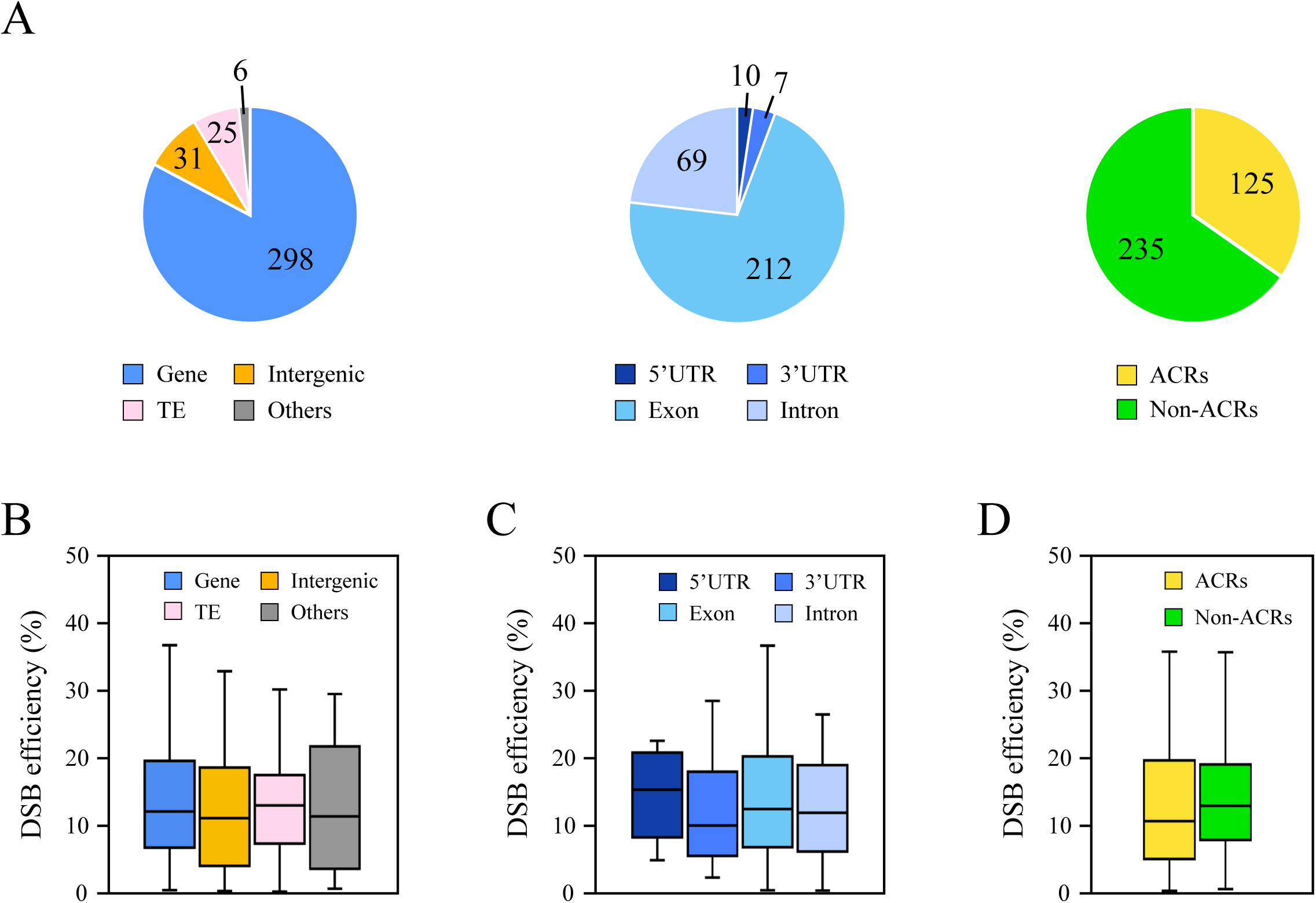
Relationship between DSB efficiency and DNA accessibility at the 360 DSB loci. (A) Classification of the 360 DSB loci based on the genetic context and the accessible chromatin regions (ACRs). First, the 360 *Sbf* I RS were classified into four groups (Gene, Intergenic, TE, and Others) based on the TAIR10 genome annotation, and then the Gene group (298 loci) was further divided into four categories (5’ and 3’ UTR, Exon, and Intron). ACRs and non-ACRs were determined in accordance with the method described in Materials and Methods. (B-D) Boxplots representing the correlation between DSB efficiency and the genetic context (B and C), or the higher-order chromatin structure (D).

### DSBs induced transient deposition of histones H4K16ac and H2A.Z around multiple cleavage sites

Studies have reported that, in human and yeast cells, both histones H4K16ac and H2A.Z are involved in DSB repair and transiently deposited on the damaged chromatin (Li and Wang 2017, González-Bermúdez et al. 2022, Gursoy-Yuzugullu et al. 2015, Xu et al. 2012); however, to the best of our knowledge, no such report has yet been published on plant cells. To evaluate the evolutionally conservation of such chromatin remodeling mechanism/dynamics via DSB-repair in plants, we performed ChIP-seq analyses of *Sbf* I-7 plants before and after HS treatment with each antibody against H4K16ac and H2A.Z (Fig. 4A), and presented H4K16ac and H2A.Z profiles 1 kb upstream and downstream at 360 DSB loci (Fig. 4B). We analyzed the data with particular cares on following two points; (1) to obtain “genuine” DSB-dependent/prone deposition of histones; (2) to discriminate those from HS-induced ones. First, we found that occupancies of both H4K16ac and H2A.Z increased over only approximately 500 bp around the DSBs compared with the level at random positions, and the majority of them were restored to their steady-state levels after 52 hours (Fig. 4B). The occupancy of these histone marks increased locally in the vicinity of most of the 360 DSB loci; however, the extent of such histone deposition continued up to 1 kb and did not spread over a wide range (Fig. 4B, C, Supplementary Fig. S4A). Second, deposition of neither H4K16ac nor H2A.Z was caused by HS treatment but was instead due to *Sbf* I-based DNA cleavage (Fig. 4B, Supplementary Fig. S4B). These increases in the levels of both histone marks were not observed in WT plants under the same condition; therefore, the majority of observed chromatin remodeling in *Sbf* I-7 plants was considered HS-independent. In addition, although we observed the peaks of H2A.Z around several dozen DSB loci before HS treatment, such H2A.Z levels after HS represented an overall increasing trend even when the DSB loci with pre-existing peaks were removed (Fig. 4B, Supplementary Fig. S4C). Altogether, these results indicated that DSBs induced H4K16ac and H2A.Z recruitment into cleavage sites in the *Arabidopsis* genome.

**Fig. 4.**
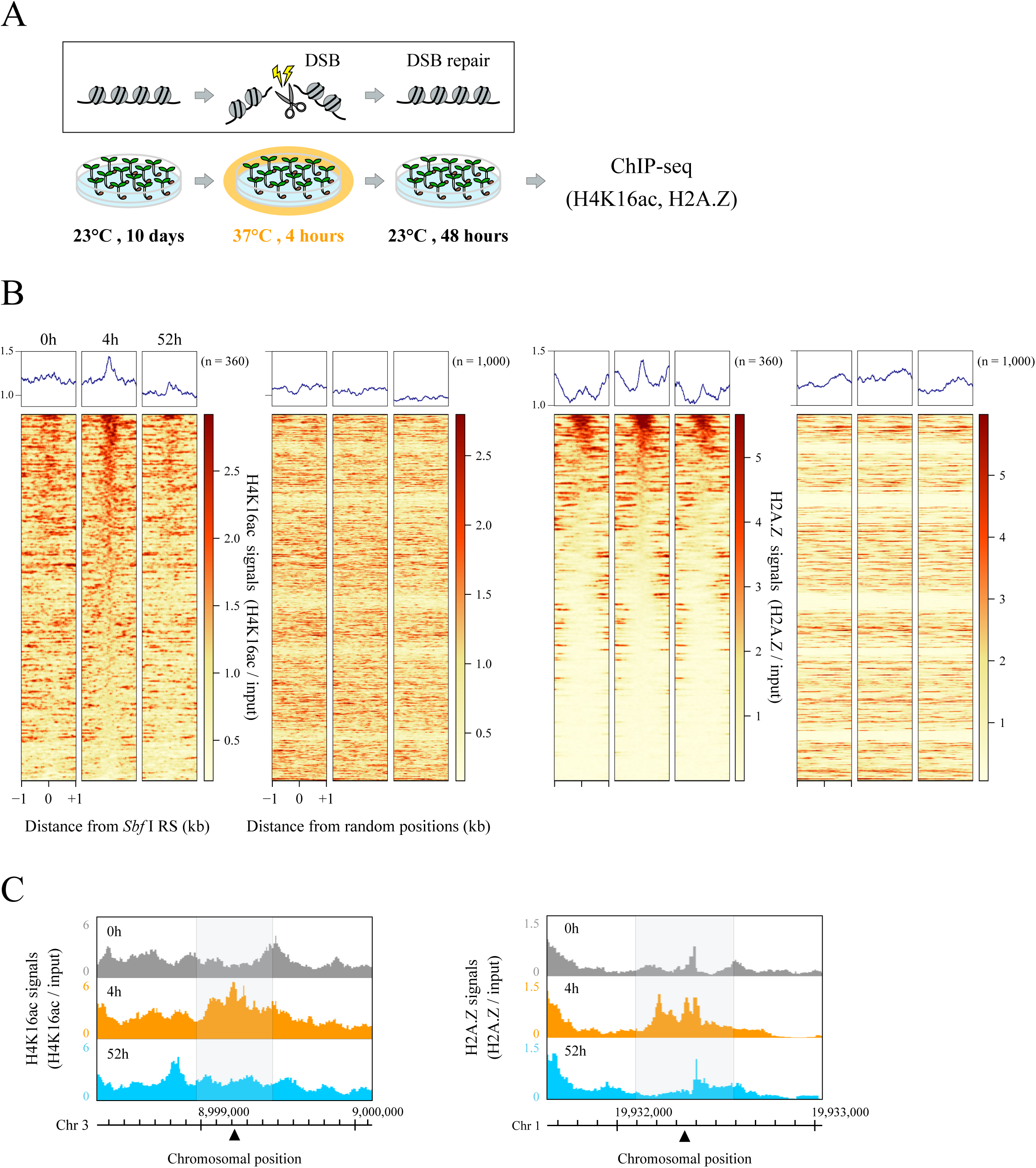
Distributions of histones H4K16ac and H2A.Z before and after DSB induction around the 360 DSB loci. (A) Experimental flow chart for DSB induction followed by ChIP-seq analysis. (B) Average profiles and heatmaps showing the occupancies of H4K16ac and H2A.Z in *Sbf* I-7 plants. These graphs and figures indicate the signal intensity of each histone mark around a 2 kb window surrounding 360 DSB loci and 1,000 random positions. All ChIP-seq data were normalized by input data, and two heatmaps of the 360 DSB loci were sorted in descending order of the total coverages of 500 bp around *Sbf* I RS for a 4h sample. (C) Genome browser screenshots representing the distributions of H4K16ac and H2A.Z around the two DSB loci (located on chromosomes 1 and 3). The gray area covers 500 bp centered on a DSB locus (black triangle).

### H4K16ac and H2A.Z dynamics during and after DSB repair

To understand how often these histone marks were deposited and removed, the dynamics of H4K16ac and H2A.Z at each DSB locus were classified based on their variation patterns. First, we classified profiles of histones H4K16ac and H2A.Z distributions during HS in WT plants to omit the influence of HS treatment on the histone dynamics (Supplementary Table S6). Second, we classified DSB loci with increased or decreased levels of H4K16ac and H2A.Z in *Sbf* I-7 plants after HS treatment, and counted those that differed from the WT classification. Those that showed the same classification as the WT were defined as “unchanged” (Fig. 5A). Classification of the variation patterns of these histone marks was performed as described in Materials and Methods using the coverage within 500 bp around the 360 S*bf* I RS. As described above, we have also shown that the mapping rate within 100 bp around the *Sbf* I RS in the *Sbf* I-7 chromosomes was reduced by the DSBs (Fig. 2A, B). To test whether this effect negatively influenced the H4K16ac and H2A.Z levels, we excluded the coverage of each histone mark within 100 bp around DSBs from the calculation of these histone levels; however, the patterns of variation of these histone marks did not change significantly (Supplementary Fig. S4D, E).

**Fig. 5.**
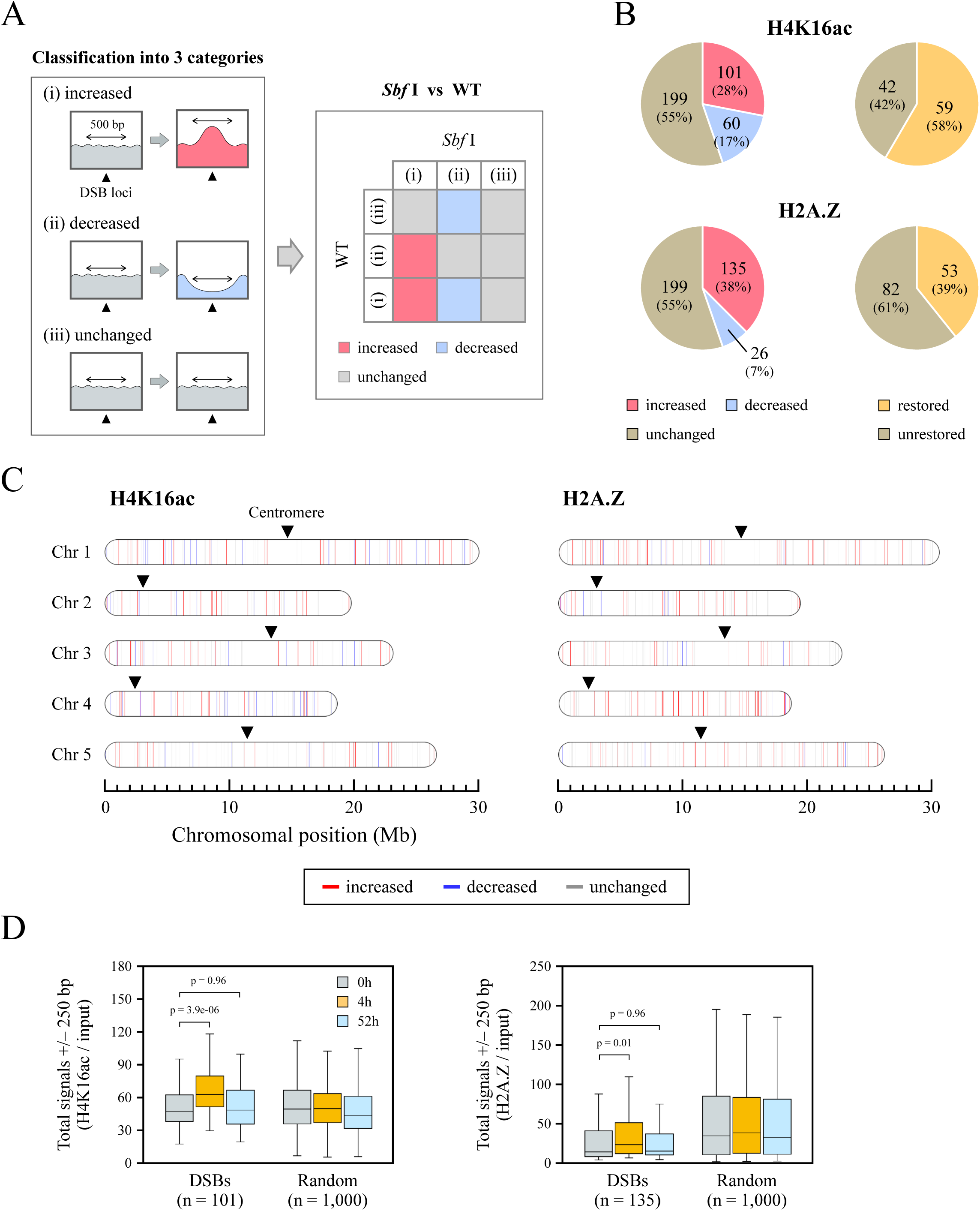
Alteration and restoration rates of H4K16ac and H2A.Z after DSB induction. (A) Framework of the classification of H4K16ac and H2A.Z distribution patterns to identify DSB-dependent chromatin remodeling. (B) Pie charts showing the rates of alteration (left) and restoration (right) of H4K16ac and H2A.Z in *Sbf* I-7 plants. (C) Alterations of H4K16ac and H2A.Z after DSB induction mapped on the *Arabidopsis* chromosomes. The colored bars indicate individual cleavage sites and corresponding alteration levels (increased: red, decreased: blue, and unchanged: gray). (D) Boxplots representing the totals of H4K16ac and H2A.Z read counts at 250 bp upstream and downstream of *Sbf* I RS or 1,000 random positions (as a negative control). Student’s t-test was used for calculation p-values (p < 0.05).

Figure 5B and Supplementary Fig. S4F show that the levels of both histones H4K16ac and H2A.Z altered at almost half of the DSB loci after HS treatment; for H4K16ac, there was an increase at 101 loci (28%) and a decrease at 60 loci (17%), whereas for H2A.Z, there was an increase at 135 loci (38%) and a decrease at 26 loci (7%). These alterations of histone occupancy were observed throughout the *Arabidopsis* genome (Fig. 5C) and, especially for increased histone occupancy, were significant overall compared with those at randomly determined positions (Fig. 5D). Under our experimental condition, the repair of cleaved DNA is expected to be completed within 48 hours (Fig. 2A, D). However, within that time, H4K16ac and H2A.Z were only completely restored to their original state at 59 of 101 (58%) and 53 of 135 sites (39%), respectively (Fig. 5B). These results may reflect a slight delay between the restoration of damaged DNA and the eviction of DSB-related histone marks. On the basis of these results, we propose that H4K16ac and H2A.Z often localize in a DSB-dependent manner and may play important roles during DSB repair.

### Co-localization of histone H4K16ac and H2A.Z after DSB induction

The incorporation of both histones H4K16ac and H2A.Z onto damaged chromatin alters the nucleosome properties and creates open, relaxed chromatin domains that extend around cleavage sites (Li et al. 2010, Xu et al. 2012, Lukas et al. 2011). Furthermore, H2A.Z exchange is essential for the subsequent acetylation and ubiquitination of the damaged chromatin (Li and Wang 2017, González-Bermúdez et al. 2022, Xu et al. 2012, Gursoy-Yuzugullu et al. 2015). Our results and these findings evoked us a hypothesis that H4K16ac and H2A.Z are co-localize on the same nucleosomes in respect to the DSB response. However, to the best of our knowledge, no such reports have been published. To verify such interaction, we integrated the profiles of H4K16ac and H2A.Z in 360 DSBs and investigated the co-localization of these histone marks by DSB induction. Interestingly, these histone marks increased together at 56 of 360 DSB loci (Fig. 6A, B). In addition, we examined the restoration rate of H4K16ac and H2A.Z at these co-localized sites after HS treatment, and found that both histone marks were restored at only 15 of the 56 sites (Fig. 6A). These results suggest that both H4K16ac and H2A.Z are often recruited at DSB loci for efficient repair but are not always co-localized and evicted simultaneously.

**Fig. 6.**
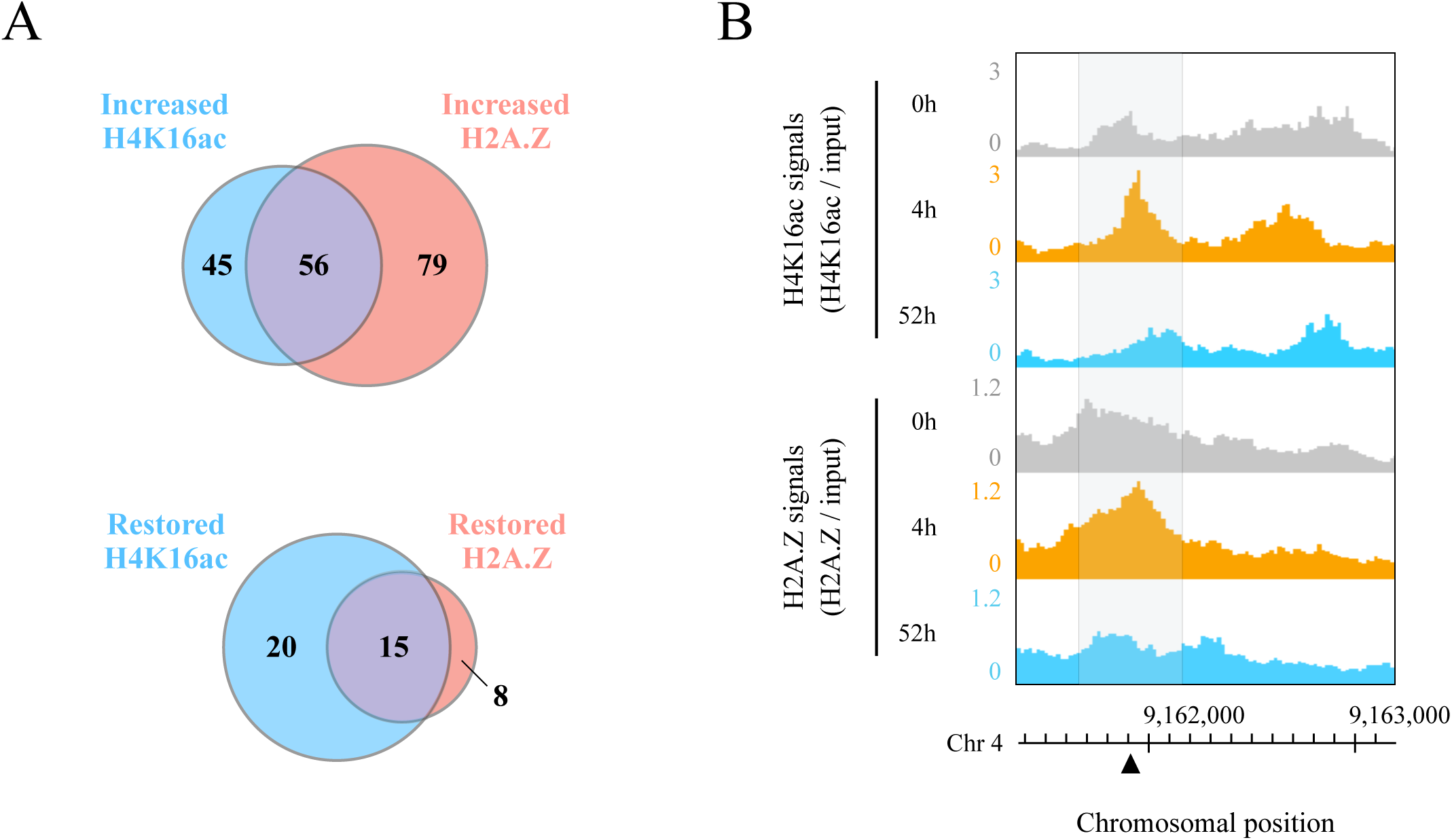
Co-localization of histone H4K16ac and H2A.Z during and after DSB repair. (A) Venn diagram of increased H4K16ac and H2A.Z peaks around 360 DSB loci, and the restoration rate of H4K16ac and H2A.Z in 56 regions where H4K16ac and H2A.Z were co-localized. (B) Genome browser screenshots showing the co-localization of H4K16ac and H2A.Z signals around a DSB locus (black triangle) located on chromosome 4. All data were normalized by input signals. Each gray area covers 500 bp centered on a DSB locus.

In summary, we established restriction enzyme-based DSB-inducible plants and showed that the DSBs induced the deposition of both histones H4K16ac and H2A.Z for accurate DSB repair in *Arabidopsis thaliana.* We also considered that such chromatin remodeling mechanism/dynamics via DSB repair is a highly conserved between plants and animals.

## Discussion

DNA double-strand breaks (DSBs) not only are a form of physical DNA damage but also affect the epigenetic landscape. To understand the mechanisms of DSB-related chromatin remodeling in detail, comprehensive and quantitative experiments must be performed that combine genome-wide chromatin analysis with a DSB induction system at various genomic loci. In this study, we established *Sbf* I heat-inducible plants in which transient DSBs can simultaneously occur at various positions in the *Arabidopsis* genome (Fig. 1). In addition, the combination of high-resolution mapping of DSBs and ChIP-seq analysis enabled us to quantitatively analyze the patterns of chromatin remodeling before and after DSB induction. We actually identified 360 *Sbf* I-dependent DSBs (Fig. 2A) and elucidated the spatial and temporal distribution patterns of histones H4K16ac and H2A.Z at their cleavage sites. We found that H4K16ac and H2A.Z increased at approximately 30% and 40% of the 360 DSB loci, respectively, and such deposition was observed throughout the *Arabidopsis* genome (Fig. 5). Intriguingly, some of these DSB loci showed co-localization of H4K16ac and H2A.Z and/or a slow rate of recovery of these histone marks (Fig. 5B, 6A). On the basis of these results, we characterized the patterns and extent of dynamic changes in the distribution of H4K16ac and H2A.Z, and elucidated that these histone marks altered everywhere throughout the *Arabidopsis* genome.

In studying the dynamics of the epigenome through the DSB repair process, *Sbf* I heat-inducible plants in which DSBs can temporarily occur at specific positions throughout the plant genome are very powerful analytical tools. Previously, DSB induction systems with CRISPR-Cas9 and restriction enzymes have been established for the analysis of the yeast and human cells; however, these methods could not elucidate the distribution of histone marks on the repaired chromosomal DNA because the induction of such nucleases was achieved by chemical inducers, and thus was not reversible (Iacovoni et al. 2010, Iannelli et al. 2017, Clouaire et al. 2018). Our expression system using the HSP18.2 promoter can very easily control the expression of the target gene by changing the environmental temperature; the transcription of the target gene is activated at 37°C and deactivated at 23°C. This heat-inducible promoter is highly compatible with many restriction enzymes including *Sbf* I because there are many alternatives with optimum reaction temperature at 37°C. In addition to these properties for the promoter and restriction enzyme, the plasmid used in this study is a fully customizable vector that contains two unique restriction sites at the *Sbf* I gene sequence and promoter/terminator junction (Supplementary Fig. S5). Therefore, this plasmid can easily replace the *Sbf* I gene portion with other restriction enzyme gene sequences, which is expected to expand the DSB region for analysis. Also, we assessed the impact of DNA accessibility, which depends on the genetic context and the higher-order chromatin structures, on the DSB efficiency, and elucidated that this feature did not correlate with DNA cleavage activity by *Sbf* I, unexpectedly (Fig. 3). In this regard, we considered that the molecular size of the *Sbf* I protein is an important factor. *Sbf* I protein is composed of 323 amino acid (AA) residues, which is much smaller than other DNA-cleaving enzymes (e.g., CRISPR-Cas9: 1,368 AA residues, TALEN: 2,500-3,500 AA residues). In addition, DNA cleavage enzymes such as CRISPR-Cas9 and TALEN have been reported to be significantly affected by the higher-order epigenetic context of their target sequences (Chung et al. 2020, Chen et al. 2016). From such difference in the protein size, we speculated that the conformation of chromosomal DNA could not have had much influence on the DNA accessibility of *Sbf* I. Taken together, our DSB induction system is considered extremely versatile for the study of the method and its related effect on DNA cleavage in plants.

In this study, we observed DSBs at 360 of 623 *Sbf* I recognition sites using *Sbf* I-inducible plants, and found that most of the damaged DNA was restored within 48 hours after heat shock (HS) treatment (Fig. 2A). However, there were some regions where DSBs remained at 48 hours after HS treatment. For the reasons given below, we ascribed this slow DSB repair to differences among cells and genomic loci in both the timing of DSB formation and the rate of DSB repair in each cell and cleavage site. The expression level and activity of introduced restriction enzymes in the plant cells vary in different tissues and developmental stages. Some cells may have high levels of both, while others may have low levels. Thus, cells with high cleavage efficiency require more time for DSB repair. Furthermore, plants and other eukaryotes have two major DSB repair pathways, which are known as homologous recombination (HR) and non-homologous end joining (NHEJ). These pathways differ in the way in which the two DNA ends are joined and the time taken for them to be completely repaired (Jasin and Rothstein 2013, Davis and Chen 2013, Kozak et al. 2009). Therefore, we also considered that the difference in the repair rate at each DSB locus observed in our analysis is due to these repair pathway choices. To resolve these issues, single-cell analysis for the quantification of DSBs and reporter assay systems that can identify the DSB repair pathway are considered to be useful.

Histone modifications and variants not only act independently but also interact with each other in a cooperative manner (Suganuma and Workman 2008, Li et al. 2017, Lu et al. 2015). In the DNA repair process of mammalian and yeast cells, studies have reported that H4K16ac and H2A.Z are transiently deposited at DSB loci to improve repair efficiency (Li et al. 2010, Xu et al. 2012, Gursoy-Yuzugullu et al. 2015). According to studies on the mechanism depositing these histone marks in DSB regions, MOF and KAT5 (Tip60) are known to be the major enzymes catalyzing the acetylation of H4K16, and in particular, KAT5 is recruited to the genomic locus where the DSB occurred (Li et al. 2017). Meanwhile, H2A.Z exchange onto DSB loci is mediated by the NuA4 complex, including Tip60 and p400 subunits (Xu et al. 2012). Notably, the NuA4 complex can deposit both H4K16ac and H2A.Z. In this study, H4K16ac and H2A.Z levels around DSBs were increased at almost half of the 360 DSB loci after HS treatment (Fig. 4), and increases in both histones occurred simultaneously at 56 DSB loci (Fig. 6). These results indicate that the DSB-dependent localization of these histone marks is a common event, at least among eukaryotes, including plants, and imply that both H4K16ac and H2A.Z are deposited via the NuA4 complex. However, whether both H4K16ac and H2A.Z are really contained in single nucleosomes remains unclear because, in this study, we only examined the localization patterns of these histone marks independently. As the solution to this problem, co-immunoprecipitation (Co-IP) experiment for elucidating the dynamics of these histone marks at more time points and identification of reader/writer proteins that participate in H4K16 acetylation and H2A.Z incorporation are necessary.

In conclusion, we introduced a heat-inducible DNA cleavage system into *Arabidopsis thaliana* that generates a sufficient number of endogenous, sequence-specific DSBs to analyze the DSB-induced chromatin landscape. This tool allowed us to analyze DSB-associated mechanisms at several positions over time and we actually discovered the new biological aspects that H4K16ac and H2A.Z localized or co-localized at DSB loci in plants. Restriction enzyme-inducible plants represent powerful tools for monitoring chromatin remodeling during DSB repair, so that events that take place under different chromatin contexts can be investigated at an unprecedentedly large scale and statistically.

## Materials and Methods

### Plant materials and growth conditions

*Arabidopsis thaliana* (Columbia ecotype) was grown at 23°C with continuous light (30-50 µmol m^−2^ s^−1^). For the genetic transformation of *Arabidopsis*, HSP18.2 promoter (Yoshida et al. 1995), SV40 nuclear localization signal (MAPKKKRKVI), full coding region of the *Sbf* I gene, and the HSP terminator sequence (Dansako et al. 2003) were subcloned into the pGreenII-MH2 vector (Hellens et al. 2000, Hirashima et al. 2006). In the following order, the constructs contained the HSP18.2 promoter, the SV40 nuclear localization signal, the S*bf* I coding region, and the HSP terminator. The resulting plasmids were introduced into *Agrobacterium tumefaciens* and the *Arabidopsis* WT plant background was transformed. Among the T1 seeds obtained, we screened the homozygous T4 plants and used them in this study.

### Methods for heat shock (HS) treatment of plants

*Sbf* I-inducible plants were stratified at 4°C in the dark for 2 days, then grown on MS medium under continuous light (30-50 µmol m^−2^ s^−1^) at 23°C for 10 days. These plants underwent HS treatment as whole plants in an NK-type artificial weather chamber (Nippon Medical and Chemical Instruments Co., Ltd.). In accordance with the procedures of Yoshida et al. (1995), Tsukaya et al. (1993), and Takahashi et al. (1992), the optimum temperature and time to promote transcription induction of the HSP18.2 promoter were 37°C and 4 hours. Therefore, HS treatment was performed under the same conditions in this study. HS-treated plants were subjected to *Sbf* I gene expression and ChIP analysis.

### RNA isolation and gene expression analysis

Total RNA was extracted from the approximately 20 individual seedlings of 10-day-old plants with the RNeasy Plant Mini Kit (QIAGEN) and rDNase (Macherey-Nagel) and reverse-transcribed with ReverTra Ace® (TOYOBO) and random hexamer. The resultant cDNA samples were used as templates for the expression analysis by quantitative RT-PCR (RT-qPCR). RT-qPCR reactions were performed using the Thunderbird® SYBR qPCR mix (TOYOBO) and Eco Real-Time PCR System (Illumina). The PCR primers used are listed in Supplementary Table S8. Relative RNA levels of the *Sbf* I gene were calculated and normalized to internal controls, the *AT5G60390* (*EF-la.*) gene-encoded GTP-binding elongation factor and the *AT4G05320* (*UBQ10*) gene-encoded polyubiquitin protein (Czechowski et al. 2005, Udvardi et al. 2008).

### Detection of DNA double-strand breaks

Genomic DNA was extracted from the same plant samples derived from RNA extraction using the DNeasy Plant Mini Kit (QIAGEN) and 0.2 ng of total DNA was used as template DNA. qPCR analysis was performed with primer sets designed to span the *Sbf* I recognition sites shown in Supplementary Table S8. Data values represent means from more than four technical replicates. The percentage of DNA double-strand breaks (DSBs) at each cleavage site was determined using the PCR amplification of intact DNA from the untreated samples as the standard, and the difference between the HS-treated samples (HS+) and the untreated samples (HS-) was calculated as the percentage of DSBs remaining. The detailed calculation method is shown below.

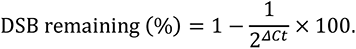

*ΔLCt* was obtained by subtraction of the threshold cycle Ct in HS-samples from the Ct in HS+ samples: *ΔLCt* = Ct (HS-) - Ct (HS+).

### Chromatin immunoprecipitation sequencing (ChIP-seq) library preparation

Approximately 150 individual seedlings of 10-day-old plants (1.0 g) were used for chromatin fixation, isolation, and fragmentation, followed by a ChIP experiment as described previously (Kudo et al. 2021, Hata et al. 2021, Zhao et al. 2020), with the following modifications. Chromatin was sonicated to 50-500 bp with a peak of 200 bp using a UD-201 ultrasonic disruptor (TOMY), with the following conditions: Output 2, Duty cycle 60%, 20 s × 18 times (interval 1 min). ChIP experiments were performed basically in accordance with the above procedure using 1 µg of an anti-H4K16ac antibody (Millipore, 07-329) and 2.4 µg of an anti-H2A.Z antibody (Kudo et al. 2021), respectively. The ChIP DNA obtained was purified using a QIAquick PCR Purification Kit (QIAGEN). We performed ChIP-qPCR to test whether the ChIP experiments were successful. ChIP-qPCR revealed that the two antibodies efficiently immunoprecipitated chromatin (Supplementary Fig. S3A). ChIP-seq libraries were prepared with sheared DNA (0.5 ng of ChIP DNA or 5.0 ng of input DNA) using ThruPLEX® DNA-seq Kit (Clontech). These obtained libraries were size-selected at 300-700 bp using ×1.0 AMPure XP beads (Beckman Coulter). Next-generation sequencing was performed using a 151 bp paired-end protocol on an Illumina HiseqX Ten platform.

### ChIP-seq data processing

The quality of each raw sequencing file is shown in Supplementary Table S7. Adapter sequences and low-quality reads were removed by Trimmomatic with the following parameters: CROP:100 ILLUMINACLIP:2:30:10 LEADING:20 TRAILING:20 SLIDINGWINDOW:4:15 MINLEN:50 (ver. 0.39; Bolger et al. 2014). To equalize the total reads of each set of ChIP-seq data, seqtk (ver. 1.3; https://github.com/lh3/seqtk) was used. All files were aligned to the reference *A. thaliana* genome (TAIR10; https://www.arabidopsis.org/) and processed using a classical ChIP-seq pipeline: Bowtie2 (ver. 2.4.5; Langmead and Salzberg 2012) for mapping and SAMtools (ver. 1.6; Li et al. 2009) for the removal of PCR-based duplicated reads (rmdup). Coverage calculations and normalization with input samples for each aligned ChIP-seq dataset (BAM files) were performed in deeptools (ver. 3.5.1; Ramírez et al. 2014). Coverage data were exported in bigwig file format for further processing. The reproducibility of each set of ChIP-seq data is shown in Supplementary Fig. S3B.

### Comprehensive analysis for genome-wide DSB detection using ChIP-seq input data

ChIP-seq data from the input DNA (0, 4, and 52h samples) were merged with three biological replicates before genome-wide DSB analysis. Using deeptools (computeMatrix tool), we calculated the total coverages of the upstream and downstream 50 bp from *Sbf* I RS. Final DSB efficiency was calculated using the following formula:

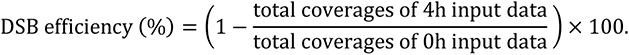

If the above calculated value was greater than 0, we judged that cleavage had occurred.

### Determination of random positions

To generate a negative control set of non-DSB regions, we first extracted 10,000 random positions on the *Arabidopsis* genome (excluding chloroplasts and mitochondria) using Perl. These random sites were filtered for being at least 10 kb away from the *Sbf* I RS. Finally, we obtained 1,000 random positions from them (Supplementary Table S3).

### Methods for extraction of accessible chromatin regions (ACRs) and classification of DSB loci in the plant genome

Based on *Arabidopsis* ACRs as defined by Ding et al. (2022), we extracted 105,301 ACRs using ATAC-seq (SRR6410823, SRR6410824), DNase-seq (SRR11144416, SRR11144417), FAIRE-seq (SRR11144419, SRR11144420), and MNase-seq (SRR10753559-SRR10753562) data from two biological replicates on PlantCADB (https://bioinfor.nefu.edu.cn/PlantCADB/). After integrating these ACRs and *Sbf* I RS information (Supplementary Table S2), 360 DSB loci were classified as 125 ACRs or 235 non-ACRs (Supplementary Table S4).

### Classification of DSB-dependent alterations for H4K16ac and H2A.Z ChIP-seq data

ChIP-seq data for each sample were analyzed by two biological replicates, and their respective genome-wide profiles were entirely consistent (Supplementary Fig. S3B). Because the ChIP-seq data obtained from the two independently prepared biological replicates showed similar profiles, each sequencing data from the two biological replicates were merged and subjected to ChIP-seq data processing. Using deeptools (computeMatrix tool), we calculated the total coverages of the upstream and downstream 250 bp from *Sbf* I RS (Supplementary Table S5). The alteration patterns of H4K16ac and H2A.Z distributions after DSB induction were classified as follows:

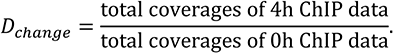

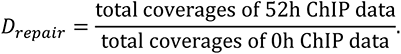

Increased, decreased and unchanged histone marks were regarded as *D_change_* ≧ 1.05, *D_change_* ≦ 0.95, and the interval between them, respectively. In addition, when *D_repair_* ≦ 1.0, these histone marks were considered restored after DSB repair; otherwise, they were considered unrestored. Similarly, we also performed calculations on WT ChIP-seq data and the DSB loci that showed the same classification as *Sbf* I-7 plants were reclassified as unchanged (Fig. 5A, Supplementary Table S6).

### Profiling methods for H4K16ac and H2A.Z ChIP-seq data before and after DSB induction

Averaged profiles and heatmaps of H4K16ac and H2A.Z ChIP-seq data were created by deeptools (plotProfile or plotHeatmap tool). The x-axis shows genomic position relative to *Sbf* I RS and the y-axis represents the mean coverage at each window (10 bp). All signals of H4K16ac and H2A.Z were normalized by input signals using deepTools (bamCompare tool). In each box-plot representation, the central line represents the median, box ends represent the first and third quartiles, and whiskers represent the minimum and maximum values without outliers. To tests differences in distribution between two populations, statistical hypothesis testing was performed using Student’s t-test. Statistical analysis was performed using R 4.1.3 with a significance level of 5% (p < 0.05).

## Data Availability

The data underlying this article are available in the article and in its online supplementary material. ChIP-seq data have been deposited to the DNA Data Bank of Japan (DDBJ) database under the accession codes DRA015426 (DRR427324-DRR427350).

## Acknowledgements

We thank A. Fukushima, K. Mukae, K. Sato, Y. Shigematsu, S. Morita, and H. Narukawa for discussion and encouragement. We also thank Edanz (https://jp.edanz.com/ac) for editing a draft of this manuscript. This work was supported by Japan Society for the Promotion of Science (JSPS) KAKENHI (22K06342 to S.S.) and Kyoto Prefectural University Academic Promotion Fund Research Encouragement Program (SK20221024 to K.K.).

## Supplementary Tables

Table S1: Information of the endogenous Sbf I recognition sites (Sbf I RS) in the A. thaliana genome.

Table S2: DSB efficiency at 360 Sbf I RS in the Arabidopsis nuclear genome.

Table S3: Information of 1,000 random positions in the *Arabidopsis* nuclear genome.

Table S4: The locations of chromatin accessible regions (ACRs) on the Arabidopsis genome and classification of Sbf I RS based on them.

Table S5: The total coverages of H4K16ac and H2A.Z at 500 bp around *Sbf* I RS before and after DSB induction.

Table S6: Determination of DSB-dependent changes in H4K16ac and H2A.Z distributions.

Table S7: Total read counts and mapping statistics of H4K16ac and H2A.Z ChIP-seq data.

Table S8: PCR primers used in study.

---

**Supplementary Fig. S1.**
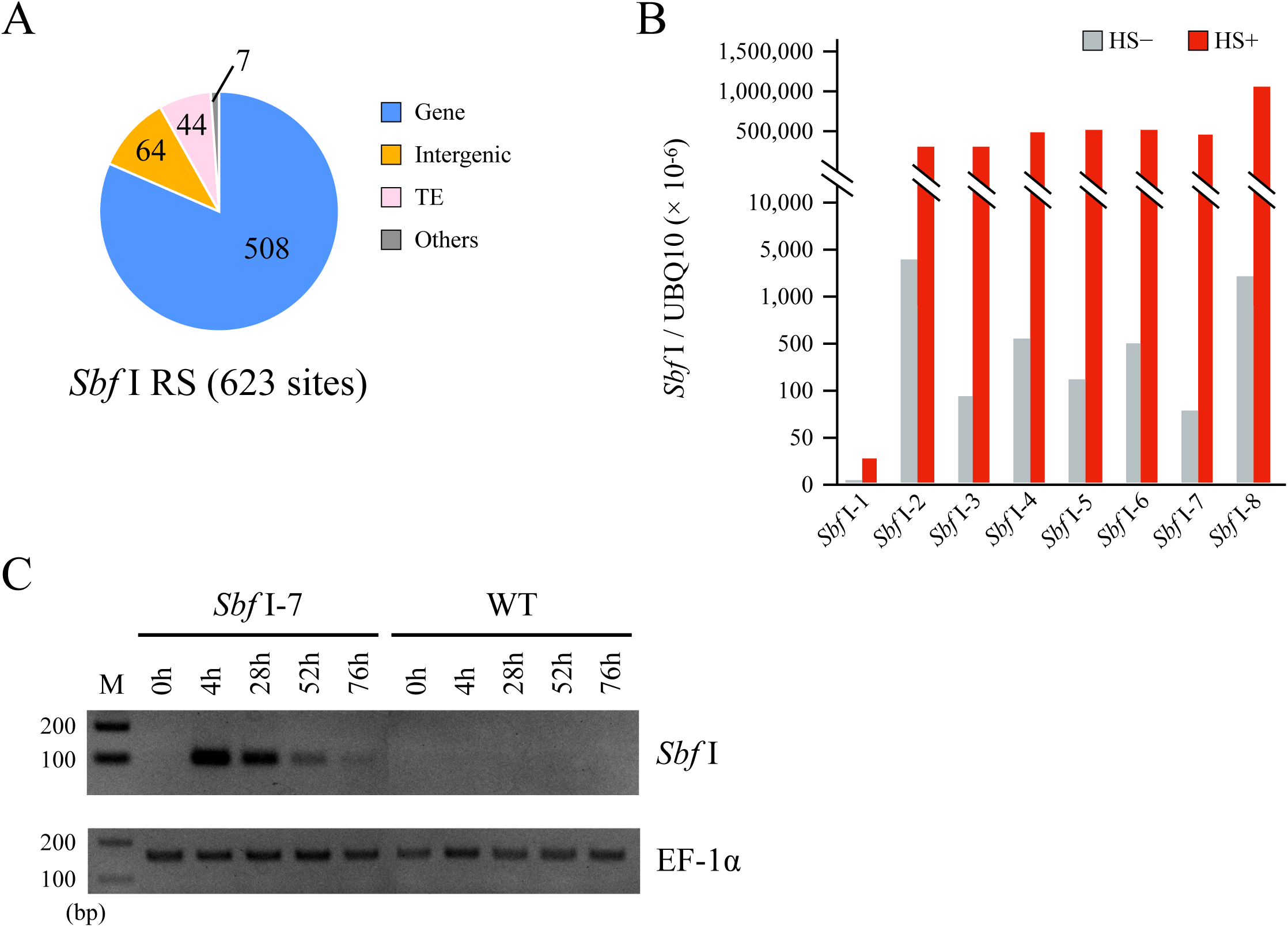
Additional information and *Sbf* I expression data in *Sbf* I-inducible plants. (A) The numbers of the endogenous *Sbf* I recognition sites (*Sbf* I RS) in the *Arabidopsis* nuclear genome. (B) Transcription level of *Sbf* I gene before and after heat shock (HS) treatment. Each value is shown the average of two independent experiments. (C) Validation of *Sbf* I gene expression by agarose gel electrophoresis. EF-1α was a control gene.

**Supplementary Fig. S2.**
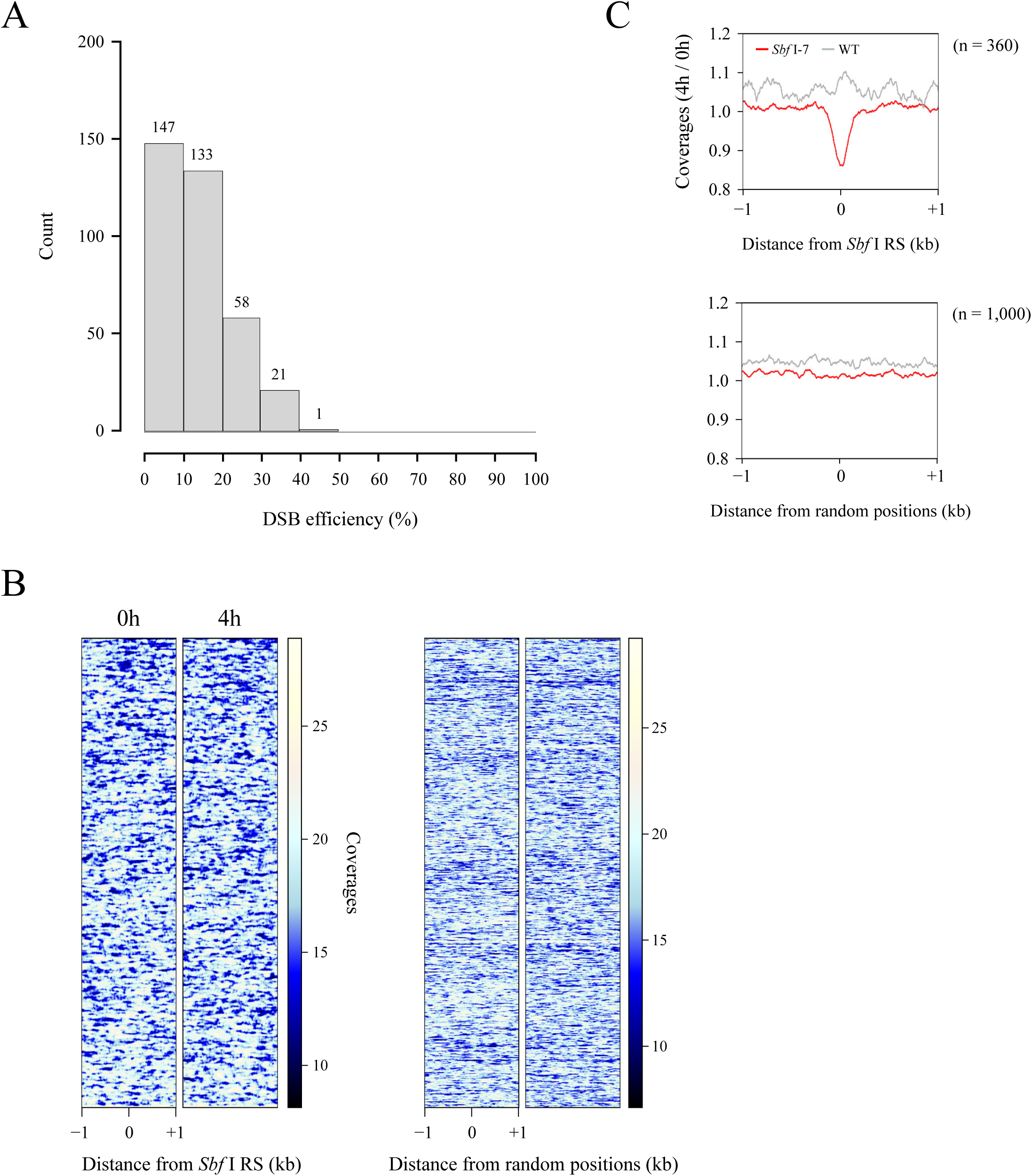
The extent of DSB efficiency at the 360 *Sbf* I RS. (A) Histogram showing the relationship between DSB efficiency and number of DSBs at the 360 *Sbf* I RS. (B) Heatmap obtained from NGS data of input DNA in WT plants. These figures represent the coverages around a 2 kb window surrounding the 360 *Sbf* I RS and 1,000 random positions. (C) Average profiles of the coverages in *Sbf* I-7 and WT plants around a 2 kb window surrounding the 360 *Sbf* I RS and 1,000 random positions. After HS treatment, DSB-dependent depletions of the coverages were observed in *Sbf* I-7 plants.

**Supplementary Fig. S3.**
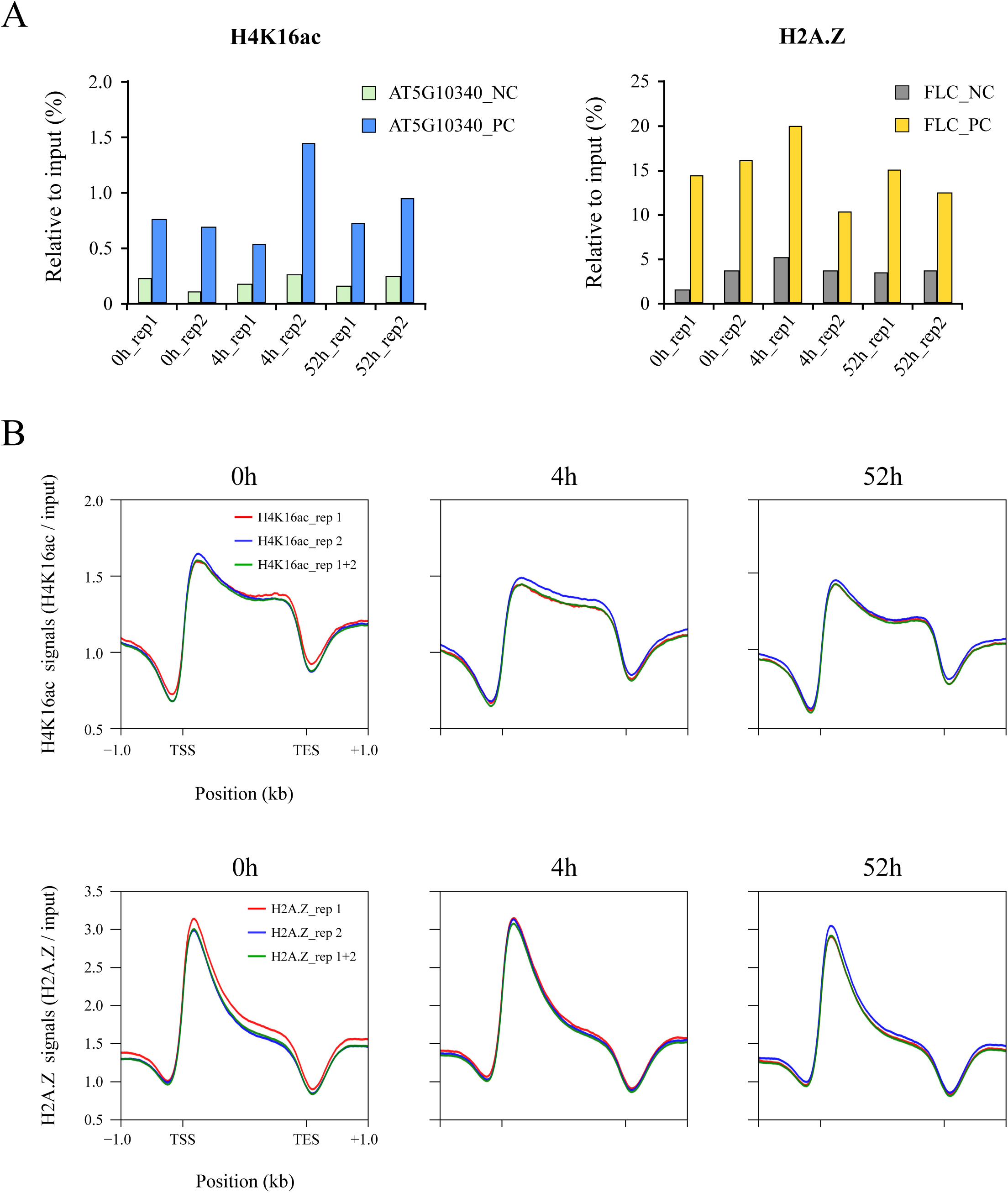
ChIP-seq data validation of histones H4K16ac and H2A.Z in *Sbf* I-7 plants. (A) ChIP experiments were conducted twice independently using two biological replicates with H4K16ac or H2A.Z antibodies. Using the primer sets shown in Supplementary Table S8, ChIP-qPCR analyses were performed at Flowering Locus C gene (*AT5G10140*) for H2A.Z enrichment check (Deal et al. 2007) or *AT5G10340* for H4K16ac enrichment check. Data values represent averages from at least two independent experiments. (B) Average profiles of histones H4K16ac and H2A.Z distributions in *Sbf* I-7 plants over *A. thaliana* genes (n= 31,109). In particular, these data were consistent with ChIP-seq analyses of histones H4K16ac and H2A.Z as shown in Wang et al. (2020) and Lu et al. (2015).

**Supplementary Fig. S4.**
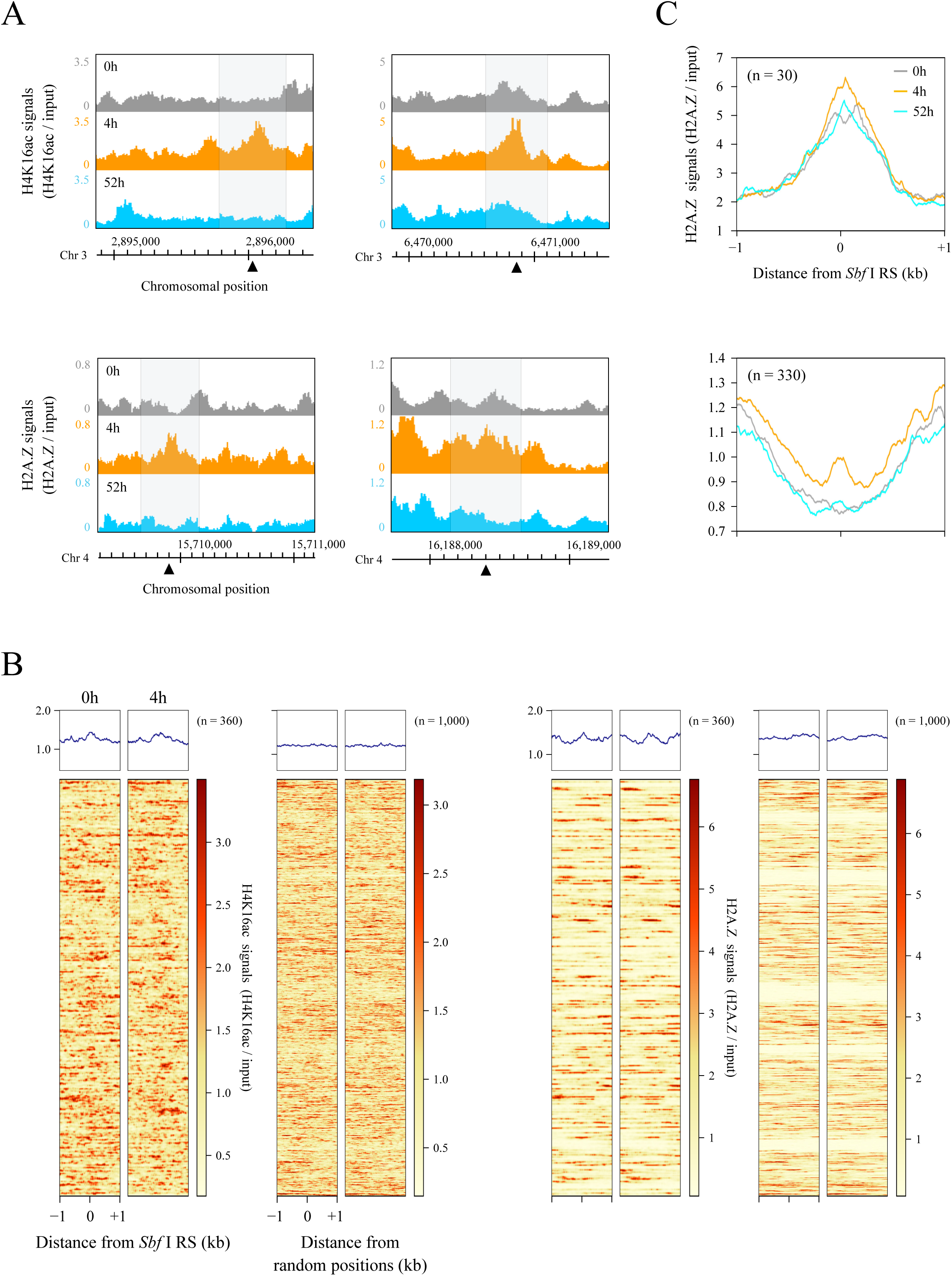
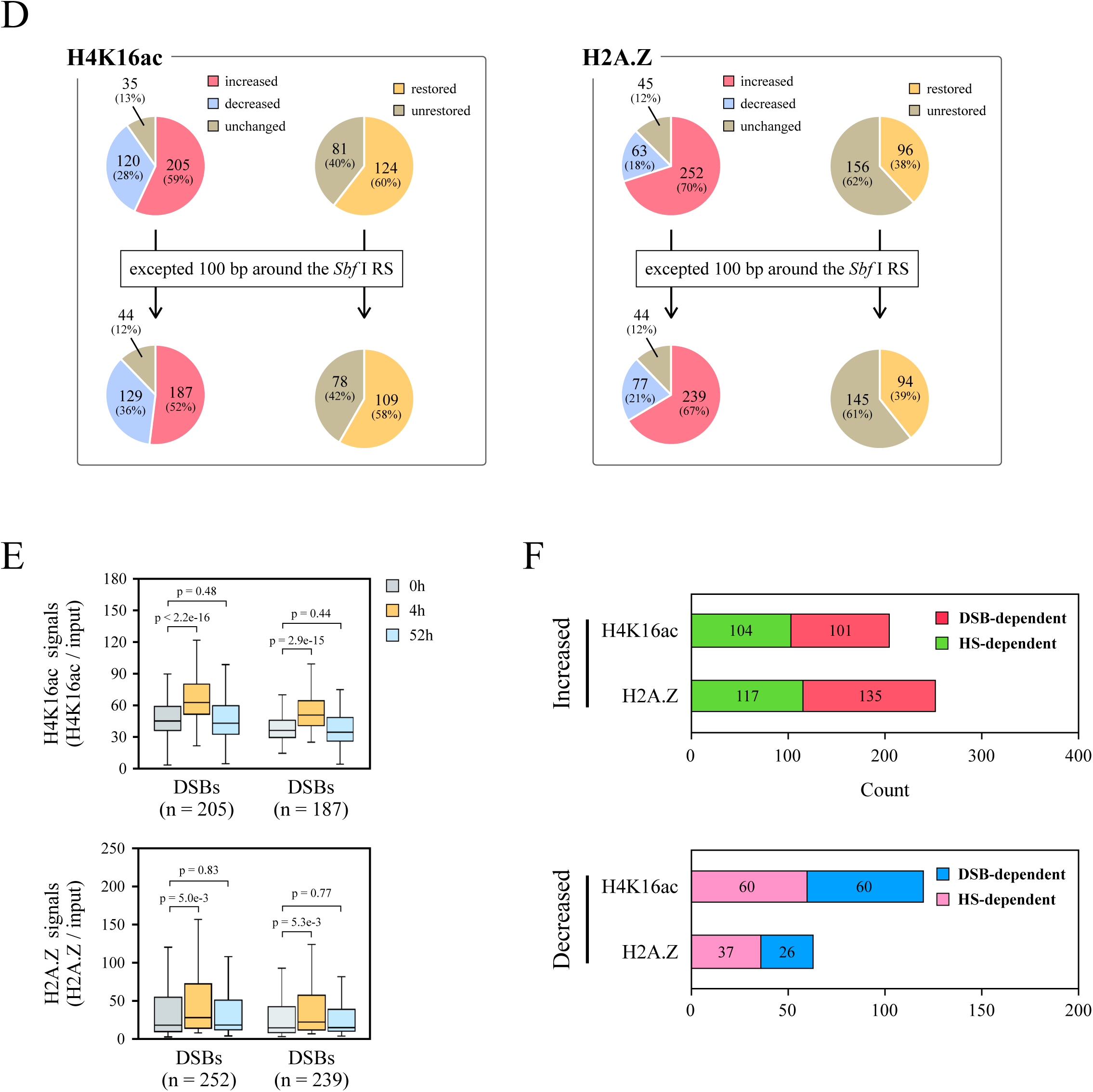
Validations of DSB-dependent alterations of H4K16ac and H2A.Z in *Sbf* I-7 plants. (A) Genome browser screenshots representing the distributions of H4K16ac and H2A.Z around other DSB loci (located on chromosome 3 and 4). All data were normalized by input signals. The gray area covers 500 bp centered on a DSB locus (black triangle). (B) Heatmaps showing the occupancies of H4K16ac and H2A.Z in WT plants (0h and 4h samples). These figures indicate the signal intensity of each histone mark around a 2 kb window surrounding 360 DSB loci and 1,000 random positions. All data were normalized by input signals. (C) Average profiles of H2A.Z signal which normalized by input for top 30 (upper) or non-top 30 (lower). (D) Pie charts representing the rate of alterations (left) and restorations (right) of H4K16ac and H2A.Z in *Sbf* I-7 plants. These graphs show the classification of two histone marks at 500 bp around *Sbf* I RS or excepted 50 bp upstream and downstream from the DSB loci. (E) Boxplots representing the total of H4K16ac or H2A.Z read counts surrounding 500 bp, excepted upstream and downstream 50 bp from *Sbf* I RS. Student’s t-test was used for calculation p-values (p < 0.05). (F) The number of cleavage sites with DSB-dependent or HS-dependent alterations of H4K16ac and H2A.Z in *Sbf* I-7 plants.

**Supplementary Fig. S5.**
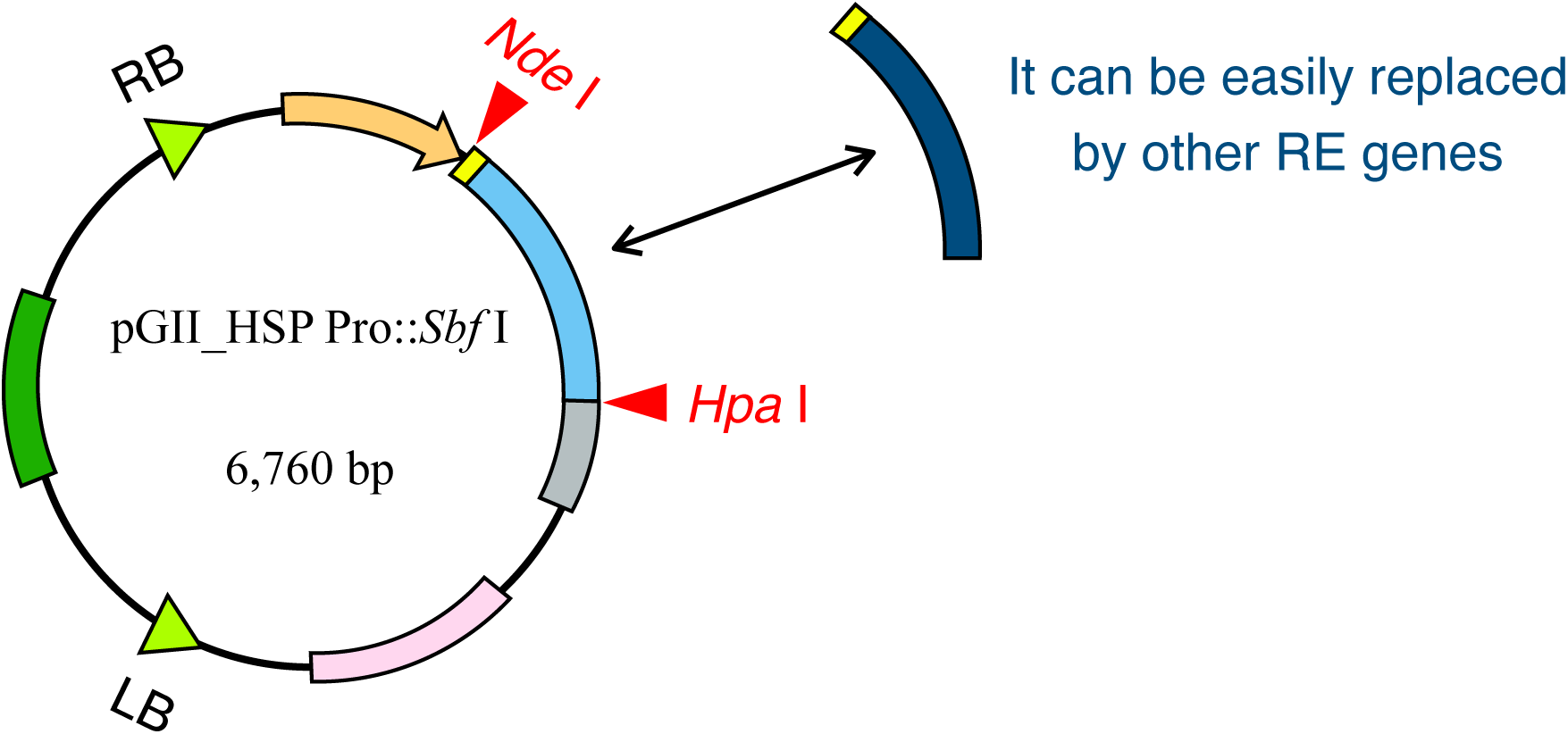
Overview and application of the improved pGreenII vector (pGII_HSP18.2 Pro::*Sbf* I) used in this study. Two unique restriction endonuclease sites (*Nde* I or *Hpa* I) are assigned to each end of the restriction enzyme (RE) gene in the T-DNA region of the plasmid shown in the figure, and these can be used to replace other RE gene sequences.

## Notes

### Competing Interest Statement

The authors have declared no competing interest.

